# The decameric repeat (DR) of PSGL-1 functions as a basic antiviral unit in restricting HIV-1 infectivity

**DOI:** 10.1101/2025.05.14.654117

**Authors:** Yang Han, Leanna Sealey, Yajing Fu, Bryan M. Delfing, Christopher Lockhart, Linda Chilin, Sameer Tiwari, M. Saleet Jafri, Dmitri Klimov, Yuntao Wu

## Abstract

PSGL-1 (P-selectin glycoprotein ligand-1) is a dimeric, mucin-like surface glycoprotein that interacts with selectins on the endothelium, facilitating leukocyte tethering and rolling during transmigration. Previous studies have demonstrated that PSGL-1 acts as an HIV restriction factor, blocking HIV infectivity by hindering particle attachment to target cells. A large part of PSGL-1’s extracellular region consists of a mucin-like domain, characterized by decameric repeats (DR), which comprise 14 to 16 DR tandems. Each DR is composed of 10 consensus amino acids (-A-T/M-E-A-Q-T-T-X-P/L-A/T-) and is notably rich in O-glycosylated threonines (30%) and prolines (10%). A proposed function of the DR is to elongate the protein backbone needed for selectin binding. However, the precise role of DR in PSGL-1’s antiviral mechanisms has yet to be fully elucidated. In this study, we performed DR deletion mutagenesis and molecular modeling, systematically deleting from one DR to all DRs, and quantified their effects on PSGL-1’s antiviral functions. Here, we demonstrate that DR is necessary for PSGL-1’s antiviral activity. Deleting DR did not affect virion incorporation of PSGL-1, but diminished PSGL-1’s ability to hinder virion attachment to target cells. We also discovered that individual single DR mutants retained 3.5 to 18% of the antiviral activity exhibited by full-length PSGL-1, and the basic antiviral activity of DR is cumulative; increasing the number of DRs correlates with heightened antiviral activity. Additionally, the intrinsic anti-HIV capability of DR was found to be transferrable; inserting the DR domain into the extracellular Ig-like domain of CD2 permitted the hybrid CD2-DR molecules to partially acquire the anti-HIV properties of PSGL-1. Further molecular modeling utilizing all-atom replica exchange molecular dynamics simulations highlighted a structure-function relationship between the anti-HIV potency of DR and the elongation of the DR backbone, as measured by DR backbone dihedral angles (*φ*) and the radial distributions of peptide atom number density (*r*_*max*_). These results collectively suggest that DR serves as an essential antiviral unit within the framework of PSGL-1’s restriction mechanism against HIV.

## INTRODUCTION

PSGL-1 (P-selectin glycoprotein ligand-1), also known as SELPLG or CD162, is a dimeric mucin-like glycoprotein primarily expressed on lymphoid and myeloid cells ^1-3^. It binds to P-, L-, and E-selectin ^1^, and functions to recruit lymphocytes into inflamed tissues during inflammation ^4-6^. This recruitment involves PSGL-1 binding to selectins on vascular surface, which initiates leukocyte tethering and rolling on endothelium, facilitating migration into inflamed tissues ^7^. Structurally, PSGL-1 possesses a relatively rigid and elongated extracellular domain (EC) that can project nearly 60 nm from the cell surface ^8,9^. The EC is also heavily glycosylated ^10,11^ and has a bending rigidity measured at 100 pNnm^2 12^. This rigidity is likely important for ligand binding, cell-cell tethering, and leukocyte rolling along the endothelial glycocalyx ^8,9^. Interestingly, overexpression of PSGL-1 can inhibit antibody binding to cell surface receptors, likely due to steric hindrance from its extended structure and heavy O-glycosylation ^13^.

A significant portion of the EC consists of a mucin–like domain with 14-16 tandem decameric repeats (DR) ^14^, where each repeat contains ten amino acids with a consensus sequence (-A-T/M-E-A-Q-T-T-X-P/L-A/T-). These repeats also contain numerous O-glycosylated threonines (30%) and prolines (10%), which were proposed to elongate and strengthen the protein backbone and separate the N-terminal selectin-binding sites from the cell membrane ^10,11,15^. DR is also associated with human PSGL-1 polymorphism that has 3 alleles (A, B, and C) with 16, 15, and 14 DRs, respectively ^14^. The A allele contains 16 DRs, whereas the B variant lacks DR#2, and the C variant lacks DR#9 and 10 ^14^. Deletion of DRs impairs PSGL-1 interaction with L- and P-selectin ^15^. In addition, DR has also been suggested to directly interact with E-selectin for supporting E-selectin-dependent rolling ^15^.

Recent research has identified PSGL-1 as an HIV-1 restriction factor that blocks HIV infectivity ^16,17^. Mechanistically, during HIV assembly, PSGL-1 is incorporated into virion particles, where it inactivates HIV through steric hindrance, preventing the virus from attaching to target cells ^17,18^. Given the critical role of DR in extending PSGL-1’s protein backbone, it is expected to be critically involved in this anti-viral mechanism. However, the specific role of DR remained unknown. In this study, we performed systematic DR mutagenesis and extensive antiviral functional mapping, integrating molecular modeling to elucidate the functional contributions of DR in the antiviral activity of PSGL-1. Our studies revealed that DR is essential for PSGL-1’s antiviral activity, acting as a basic unit in restricting HIV-1 infectivity. Individual single-DR mutants retained 3.8 to 18% of the antiviral activity of the full-length PSGL-1. Molecular modeling further highlighted a structure-function relationship between the antiviral potency of DR and its backbone elongation, suggesting that DR serves as a critical component within the framework of PSGL-1’s restriction against HIV.

## MATERIALS AND METHODS

### Cells and Cell culture

For cell culture, HEK293T (ATCC) and HeLaJC.53 (kindly provided by Dr. David Kabat) were maintained in Dulbecco’s modified Eagle’s medium (DMEM) (Invitrogen) containing 10% heat-inactivated FBS (Neuromics) and 50 units/mL of penicillin and 50 µg/mL of streptomycin (Invitrogen). HIV Rev-A3R5-GFP indicator cell (kindly provided by Virongy) were cultured in RPMI-1640 plus 10% FBS supplemented with 1 mg/mL geneticin (G418) (InvivoGen) and 1 μg/mL puromycin (Gibco).

### Plasmids, vectors, transfection, virion production and purification

The infectious HIV-1 molecular clone pNL4-3 was obtained from the NIH AIDS Reagent Program. pCMV3-Empty was purchased from Sinobiological. pCMV6-XL5-CD2 was purchased from Origene. PSGL-1 DR mutants were synthesized from Twist Bioscience, including pcDNA3.1-PSGL1, pcDNA3.1-PSGL1-15DR, pcDNA3.1-PSGL1-13DR, pcDNA3.1-PSGL1-11DR, pcDNA3.1-PSGL1-9DR, pcDNA3.1-PSGL1-7DR, pcDNA3.1-PSGL1-5DR, pcDNA3.1-PSGL1-3DR, pcDNA3.1-PSGL1-2DR, and pcDNA3.1-PSGL1-ΔDR were synthesized from Twist Bioscience. The PSGL-1 single DR mutants were also synthesized from Twist Bioscience, including pcDNA3.1-PSGL1-DR#1, pcDNA3.1-PSGL1-DR#2, pcDNA3.1-PSGL1-DR#3, pcDNA3.1-PSGL1-DR#4, pcDNA3.1-PSGL1-DR#5/14, pcDNA3.1-PSGL1-DR#6, pcDNA3.1-PSGL1-DR#7/13, pcDNA3.1-PSGL1-DR#8, pcDNA3.1-PSGL1-DR#9, pcDNA3.1-PSGL1-DR#10, pcDNA3.1-PSGL1-DR#11, pcDNA3.1-PSGL1-1DR#12, pcDNA3.1-PSGL1-1DR#15 and pcDNA3.1-PSGL1-DR#16.

The APOBEC3G vector, pcDNA3.1-A3G, was synthesized using the pcDNA3.1 vector by Twist Bioscience. A pCMV6-SERINC5 vector was kindly provided by Dr. Yonghui Zheng.

The CD2 mutants with DR insertion were synthesized from Twist Bioscience, including pCMV-CD2-TriDR and pCMV-CD2-16DR. For HIV-1 virus production, HEK293T cells were cotransfected in a 6-well plate with 1 μg of HIV-1 DNA (pNL4-3) plus the indicated doses of PSGL-1, PSGL-1 mutant, or a pCMV3-Empty vector, using transfection reagent as recommended by the manufacturer (Virongy). Virion-containing supernatants were collected at 48 hours post transfection.

### Western Blots

Cells were lysed in LDS lysis buffer (Invitrogen), proteins were denatured by sonication and boiling. Samples were subjected to SDS-PAGE using Invitrogen NuPAGE 4–12% Bis-Tris gels, transferred to nitrocellulose membranes, and incubated overnight at 4°C with an anti-human CD162 Antibody (KPL-1) (BioLegend) (1:1000 dilution). Membranes were incubated with HRP-linked anti-mouse IgG antibody (Cell Signaling Technology, Cat#7076S, 1:2000 dilution), for 1 h at room temperature. For GAPDH staining, we incubated membrane at 4°C overnight with anti-GAPDH goat polyclonal antibody (Abcam) (1:500 dilution), followed with peroxidase-labeled rabbit anti-goat IgG (H+L) antibodies (Seracare) (1:2500 dilution) for 1 hour at room temperature. Chemiluminescence signal was detected by using West Femto chemiluminescence substrate (Thermo Fisher Scientific), and images were captured with a CCD camera (FluorChem 9900 Imaging Systems, Alpha Innotech).

### p24 ELISA

Detection of extracellular HIV-1 p24 was performed using an in-house p24 ELISA kit. Briefly, microtiter plates (R&D Systems) were coated with anti-HIV-1 p24 monoclonal antibody (NIH HIV Reagent Program, cat#3537). Samples were incubated for 2 hours at 37 °C, followed by washing and incubating with biotinylated anti-HIV immune globulin (Thermo) for 1 hour at 37 °C. Plates were then washed and incubated with avidin-peroxidase conjugate (R&D Systems) for 1 hour at 37°C, followed by washing and incubating with tetramethylbenzidine (TMB) substrate. Plates were kinetically read using an Agilent BioTek ELx800 Microplate Reader at 630 nm and 450 nm.

### Viral Attachment Assay

HIV-1 viral particles produced in the presence of PSGL-1 or PSGL-1 mutants were incubated with HelaJC.53 cells (prechilled at 4 °C for 1 hour) at 4°C for 2 hours. The cells were then washed extensively (5 times) with cold PBS buffer and then lysed with LDS lysis buffer (Invitrogen) for analysis with Western blot.

### Virion Incorporation of PSGL-1 and its mutants

HIV-1 particles were assembled in 6-well plates by cotransfecting Hek293T cells with 1 μg HIV-1 DNA (pNL4-3) plus 500 ng of either a PSGL-1 vector, an empty vector, or a vector expressing PSGL-1 mutant. Briefly, magnetic Dynabeads Pan Mouse IgG (Invitrogen) (1 × 10^8^ beads/0.25 mL) were conjugated with mouse anti-PSGL-1 antibody (KPL-1) (BD Pharmingen) for 30 minutes at room temperature. After conjugation, antibody-conjugated beads were incubated with viral particles for 1 hour at 37 °C. The bead-virus complexes were then pulled down with a magnet and washed with cold PBS for 5 times. Captured viral particles were eluted in 10% Triton X-100 PBS and quantified by p24 ELISA.

### Infectivity assays and quantification of half-maximal inhibitory dosage (IC_50_)

For flow cytometry-based infectivity assay, virus particles were produced as described in virion production. Rev-A3R5-GFP indicator cells were infected with virus (0.2-0.3 million cells/infection). The cells were then washed with media and cultured in 1 mL fresh media. GFP expression was analyzed by flow cytometry at 48 or 72 h post infection using a FACSCalibur system (BD Biosciences). The percentage of GFP+ cells was quantified. Half-maximal inhibitory concentration (IC_50_) was calculated by normalizing HIV-1 control as 100% and then plotted in a GraphPad Prism 9 software using a non-linear regression (curve fit).

### Molecular simulation systems

To study the effect of amino acid sequence on DR behavior, we modelled DRs with sialyl Lewis X O-glycans using the CHARMM36m protein force field ^19^ and CHARMM36 carbohydrate force field ^20^. Sialyl Lewis X O-glycan 2 was chosen based on previous structural analysis ^21^. CHARMM-modified TIP3 water ^22,23^ was used to solvate the system with 150 mM of NaCl salt, adding counterions to neutralize the system depending on the DR sequence. The size of the cubic simulation box was adjusted to create a ∼10 Å water buffer between the solute and a box edge.

### Replica-Exchange Molecular Dynamics Simulations

We employed Replica Exchange with Solute Tempering (REST) to probe the conformational ensembles of DRs. REST simulations were performed using the program NAMD ^24^. REST enhances conformational sampling by replicating a simulation system along an external condition, e.g., temperature, and periodically attempting replica exchanges between neighboring conditions according to the Metropolis criterion. REST differs from classical replica exchange simulations ^25^ by scaling solvent interactions effectively keeping the solvent cold and removing it from Metropolis criterion during exchange attempts. The benefit of this approach is that the number of replicas can be reduced thereby easing computational burden while maintaining sampling efficiency ^26^. Several variants of REST exist ^27 24,28 29^. Our REST implementation tempered protein, glycans, and counterions, which constitute a solute. Solvent, which includes water and salt, was kept “cold” due to scaling of its interactions by the factor *T*_*r*_ /*T*_0_, where *T*_*r*_ refers to the thermostat temperature for condition *r* and *T*_0_ is the target temperature for analysis ^28 26^. Interactions between solute and solvent were partially tempered with a factor of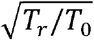. Our REST simulations utilized periodic boundary conditions. Electrostatic interactions were computed with Ewald summation and van der Waals interactions were smoothly switched off in the interval from 8 to 12 Å. Hydrogen atom covalent bonds were constrained by the ShakeH algorithm. Temperature was controlled with underdamped Langevin dynamics with a damping coefficient of 5 ps^-1^. Pressure was held constant at 1 atm using the Langevin piston method. An integration step of 1 fs was used. In total, our REST simulations for a DR employed 16 conditions with a geometric temperature distribution from 310 to 500 K. Replica exchanges were attempted every 2 ps between neighboring temperatures. Each DR system was run for a total of 120 ns per replica, yielding 1.92 μs overall per system. For each DR, sampling within the last 60 ns at 310 K was reserved for analysis (See **Supporting Information**)

### Structural Probes

We evaluated several structural quantities, which showed a significant correlation with the potency against HIV infection. First, we computed the distributions of DR backbone dihedral angles, *φ* and *ψ*, using the program VMD ^30^.

Second, we constructed the radial distribution functions *g*(*r*) measuring the number density of peptide atoms as a function of distance *r* to the peptide center of mass. Specifically, *g*(*r*) has been defined as

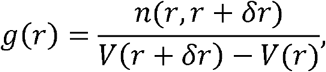

where n(*r, r* + δ*r*) is the number of peptide (excluding glycan) atoms occurring within the distances *r* and *r*+ δ*r* from the peptide center of mass and V(*r*+ δ*r*) - V(*r*) is the volume between two concentric spheres with the radii *r* and *r*+ δ*r*. To compute *g*(*r*) we applied a bin size δ*r* = 0.5 Å. From *g*(*r*), the location of number density maxima *r*_*max*_ was computed for each DR as an average of instantaneous maxima locations for individual frames. Because the dihedral angles and radial distribution functions exclude glycan atoms, they probe the conformations of peptide within DR. To understand the relation between structural quantities and DR potency against HIV-1, we computed Pearson’s correlation coefficients *ρ*. The stability of these correlations was assessed using a leave-one-out cross-validation scheme (see **Supporting Information**). All structural quantities and their correlations with the potency were evaluated at *r*_0_ = 310 K. The examples of radial distribution functions *g*(*r*) for DRs are plotted in **Fig. S4**.

### Clustering

We utilized the method described by Daura et. al ^31^ to perform density-based conformational clustering of the DR poses produced by our REST simulations. Clustering was implemented based on the pairwise RMSD distributions of peptide (excluding glycan) heavy atoms. Prior to RMSD measurement, the peptides were aligned based on their coordinates. For our clusters, we used the cutoff value of *r*_C_ = 3 Å.

Clustering was applied to 10,000 structures sampled across the equilibrated dataset (**See Supporting Information**). Clusters with a minimum population of 10% were retained for further analysis.

### Data analysis

Statistical analyses were performed with GraphPad Prism 9.5 (GraphPad Software). Paired t-tests were applied to compare two groups in the figures. Applied statistical analyses are specified in the figure legends. Data are presented as mean ± SD of three replicates unless stated otherwise in the figure legend.

## RESULT

### PSGL-1’s DR domain is required for its antiviral activity

Structurally, a large portion of PSGL-1’s extracellular domain consists of 14 to 16 heavily glycosylated, mucin-like decameric repeats (DRs) (**Fig.1A**). To determine the involvement of DR in PSGL-1’s anti-HIV activity, we performed DR deletion mutagenesis, in which the whole DR domain with all 16 DRs was deleted from PSGL-1 to create PSGL-1ΔDR. The deletion was confirmed by Western blot (**Fig. 1B**), and subsequently quantified for impacts on PSGL-1’s anti-HIV activity. HIV particles were assembled in the presence of PSGL-1 or PSGL-1ΔDR (1 *μ*g HIV DNA plus 400 ng of PSGL-1 or PSGL-1ΔDR), and virion infectivity was quantified by infecting an HIV Rev-dependent indicator T cell, Rev-A3R5-GFP ^32,33^, using an equal p24 level of virus input. At 400 ng vector dose, the full-length PSGL-1 completely inactivated HIV-1 virion infectivity, whereas deleting the DR region (PSGL-1ΔDR) largely abolished this anti-HIV activity (**Fig. 1C** and **Fig. S1**). These results demonstrated that the DR domain plays a critical role in PSGL-1’s antiviral activity.

**Fig. 1.**
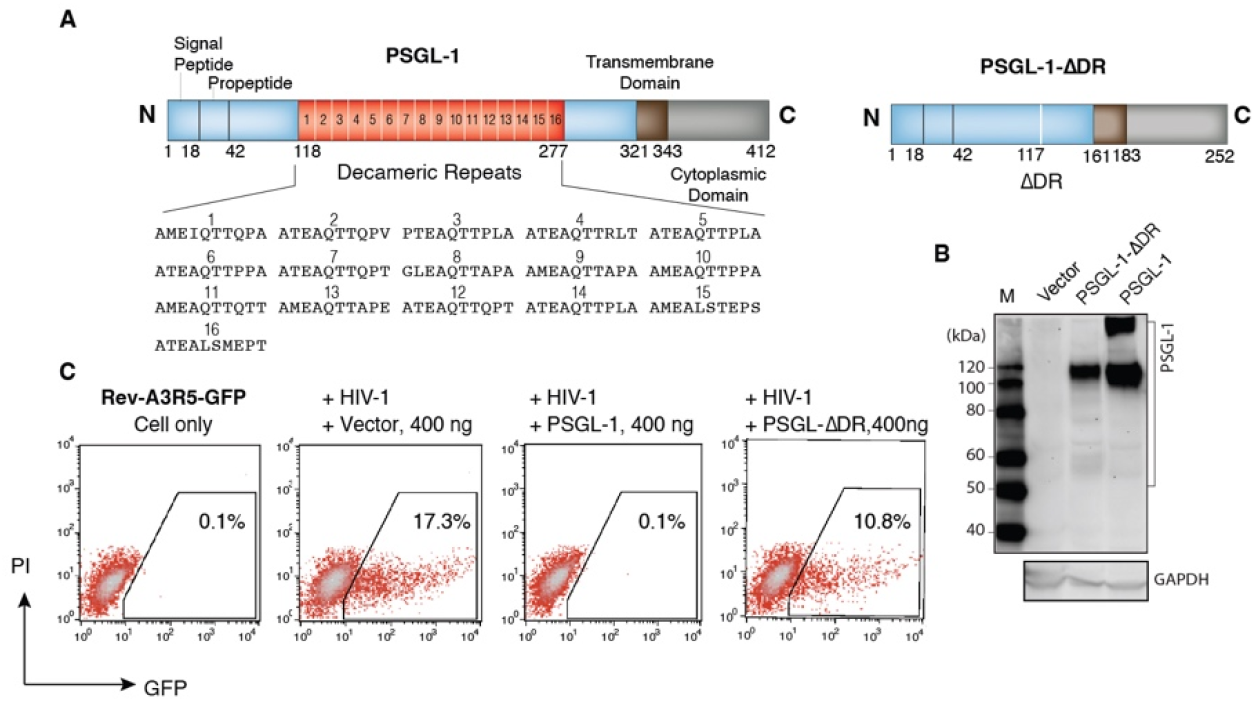
PSGL-1’s DR domain is required for its antiviral activity. **(A)** Schematic of PSGL-1 and PSGL-1ΔDR mutant. (**B**) Western blot detection of PSGL-1 DR deletion mutant protein. HEK293T cells were transfected with 400 ng DNA of PSGL-1, PSGL-1ΔDR mutant, or an empty vector. Cells were lysed and analyzed with an anti-PSGL-1 antibody to detect the PSGL-1 expression. The blot was also probed with an anti-GAPDH antibody for loading control. (**C**) Quantification of the antiviral activities of the DR deletion mutant. HEK293T cells were co-transfected with 1 μg pNL4-3 DNA plus 500 ng PSGL-1 or PSGL-1ΔDR, or an empty vector. Virions were harvested at 48 hours post-transfection and used to infect Rev-A3R5-GFP indicator cells, using an equal p24 level of virus input. Virus infectivity was quantified with flow cytometry of HIV^+^ cells (GFP^+^). PI (propidium iodide) was used to examine GFP only in viable cells. Representative result from experimental triplicate.

### The cumulative effect of DR in restricting HIV-1 infectivity

The DR domain contains 14-16 decameric repeats. Given that deleting all DRs greatly diminished PSGL-1’s antiviral activity (**Fig. 1C**), we investigated whether there is a minimal number of DRs required for PSGL-1’s activity. Thus, we performed serial DR deletion mutagenesis and created a panel of PSGL-1 mutants carrying various numbers of DRs, from 15 DRs to a single DR (**Fig. 2A**). Expression of mutant PSGL-1 proteins was confirmed by western blot (**Fig. 2B**). For an initial evaluation, virus particles carrying each individual DR mutant were assembled using 1 μg HIV DNA plus 500 ng of PSGL-1 or each individual DR mutant DNA. Virion infectivity was quantified in Rev-A3R5-GFP cells using an equal p24 level of virus input. As shown in **Fig. 2C**, for PSGL-1 mutants carrying 3 and more DRs, at the high vector dosage (500 ng), we observed a near complete inhibition of HIV-1 infectivity, similar to that of the full-length PSGL-1. For the 2 DR mutant, we observed a slight reduction of the anti-HIV activity, which is at approximately 83% of the full-length PSGL-1 (100%). Surprisingly, two of the single DR mutants, PSGL-1DR#16 and PSGL-1DR#1, also inhibited HIV at around 80-90% of the full-length PSGL-1 (**Fig. 2D**). Nevertheless, these inhibitions were observed at the high saturating dosage (500 ng).

**Fig. 2.**
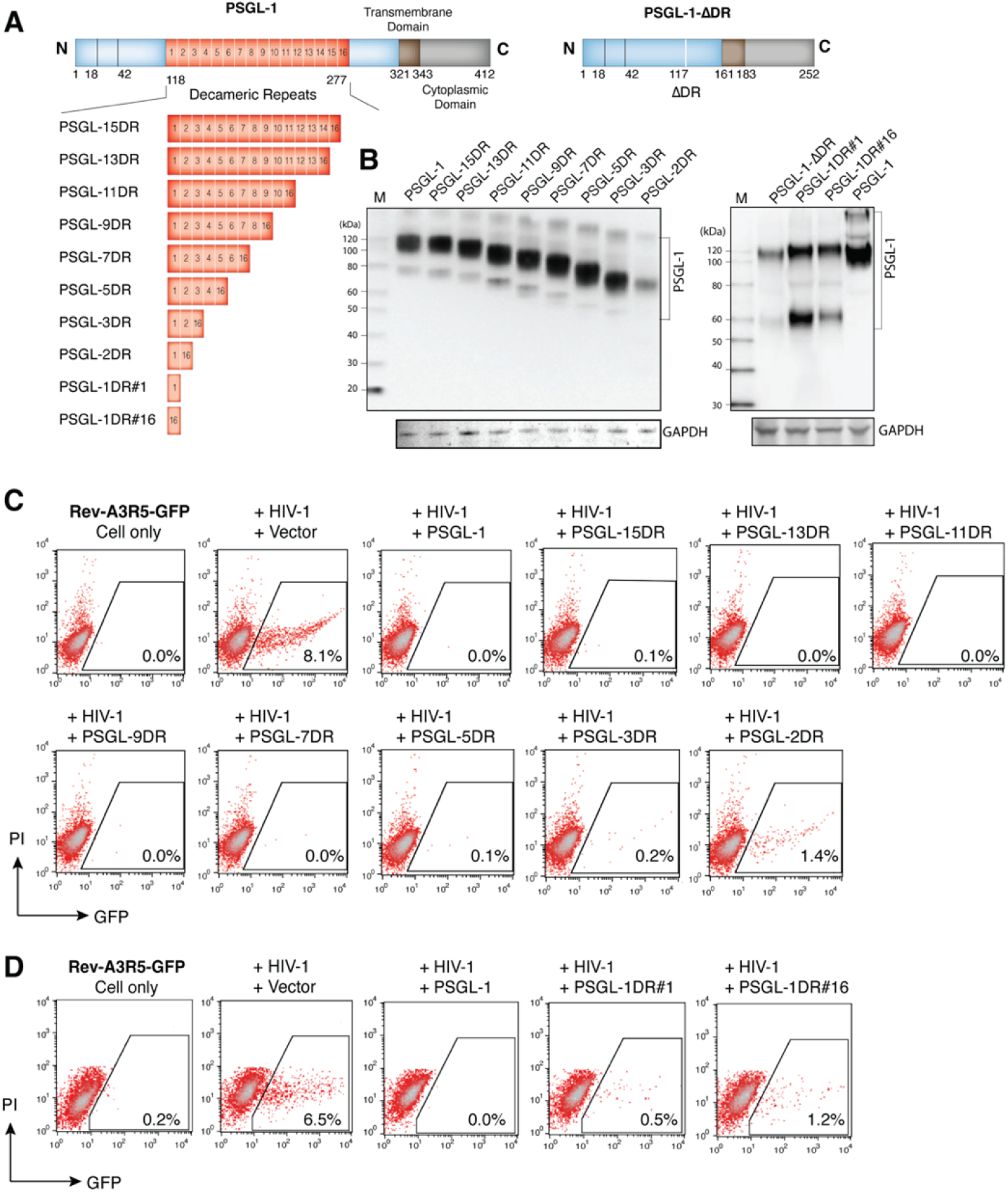
Serial DR deletion mutagenesis study. **(A)** Schematic of DR deletion mutagenesis. (**B**) Western blot detection of PSGL-1 DR deletion mutant proteins. HEK293T cells were transfected with 400 ng DNA of PSGL-1 or individual PSGL-1 DR deletion mutants. Cells were lysed and analyzed with an anti-PSGL-1 antibody to detect the expression of PSGL-1. The blot was also probed with an anti-GAPDH antibody for loading control. (**C** and **D**) Quantification of the antiviral activities of the DR deletion mutants. HEK293T cells were co-transfected with 1 μg pNL4-3 DNA plus 500 ng PSGL-1, individual PSGL-1 DR deletion mutants, or an empty vector. Virions were harvested at 48 hours post-transfection and used to infect Rev-A3R5-GFP indicator cells, using an equal p24 level of virus input. Virus infectivity was quantified with flow cytometry of HIV^+^ cells (GFP^+^) at 48 hours post infection. Representative result from experimental triplicate.

To more accurately quantify the antiviral activities of the DR mutants, we further performed PSGL-1 dosage-dependent inhibition assays to measure the IC_50_ values, particularly those of the 2 DR and 1 DR mutants. We also chose PSGL-1, PSGL-1-7DR, and PSGL-1ΔDR for comparison. Viruses were assembled in the presence of 1 to 500 ng of each DR mutant, and then quantified for virion infectivity using an equal p24 level of virus input (**Fig. 3A** to **3C** and **Fig. S2**). As shown in **Fig. 3D**, the full-length PSGL-1 had an IC_50_ of 2.5 ± 1.2 ng (experiment repeats, n = 10), whereas PSGL-1-7DR had an IC_50_ of 9.0 ± 0.9 ng (n = 5). Deleting all DRs (PSGL-1ΔDR) led to the increase of IC_50_ to greater than 400 ng (n = 3). For the PSGL-1-2DR mutant, the IC_50_ was measured at 12.6 ± 2.6 ng (n = 5), whereas for the PSGL-1-1DR mutants, the IC_50_ values varied considerably, with the IC_50_ at 26.0 ± 11.1 ng (n = 6) for PSGL-1-DR#1, and the IC_50_ at 61.0 ± 14.1 ng (n = 6) for PSGL-1-DR#16. These results demonstrated that there is a cumulative effect; increasing the number of DRs is associated with increased anti-HIV activity. Conversely, deleting the DRs to a single DR led to a 10-25 fold reduction of PSGL-1’s antiviral activity. Nevertheless, even a single DR mutant can still maintain 4-10% potency of PSGL-1 (%potency = IC_50_ PSGL-1/IC_50_ DR mutant). The IC_50_ variations of the two 1DR mutants likely result from the sequence variations of the decameric repeats (**Fig. 4A**).

**Fig. 3.**
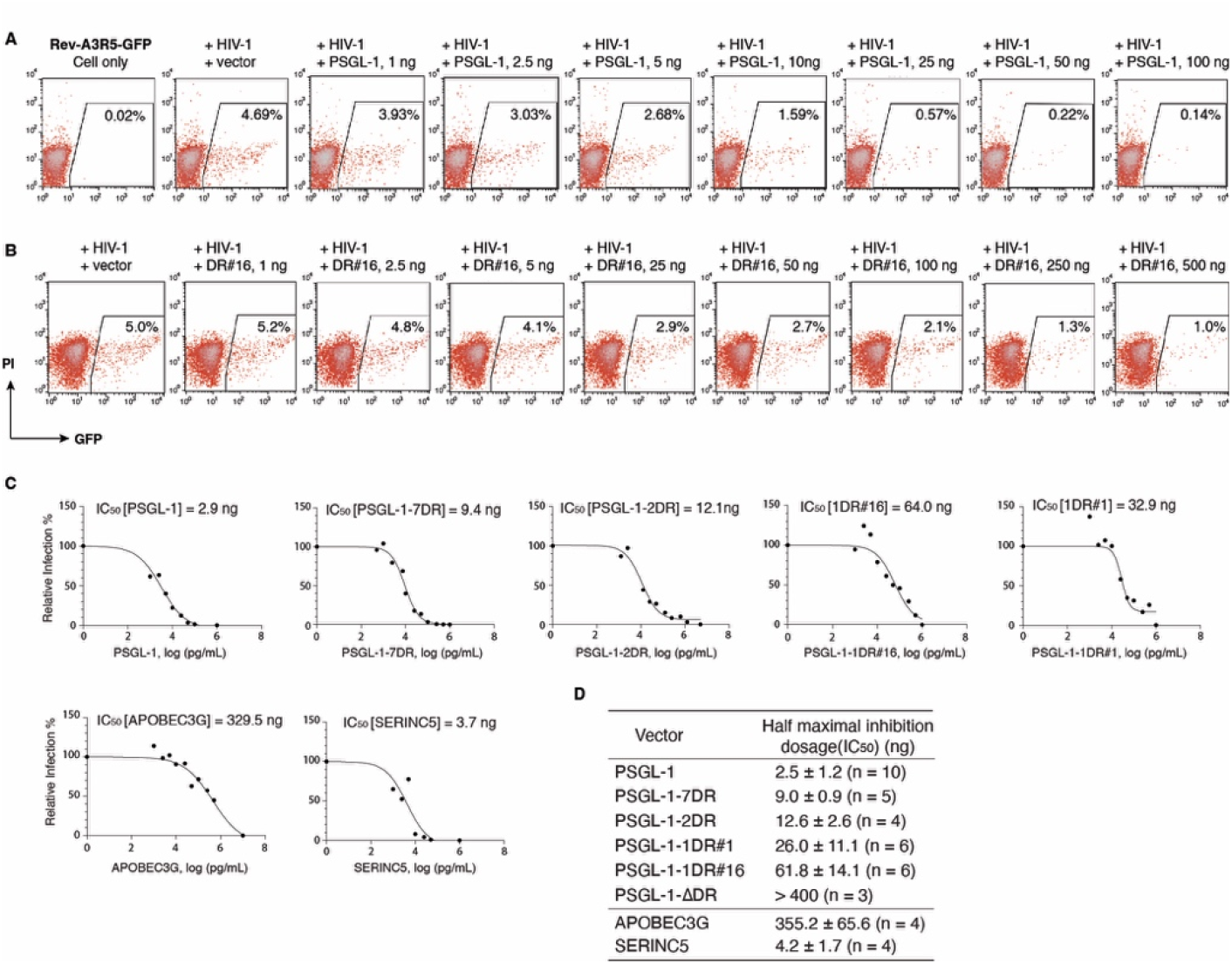
The cumulative effect of DR in restricting HIV-1 infectivity. **(A** and **B)** Accurate quantification of the 50% inhibitory dosage (IC_50_) of PSGL-1 and its DR deletion mutant (PSGL-1-DR#16) using the HIV Rev-dependent Rev-A3R5-GFP cells. HEK293T cells were cotransfected with 1 μg pNL4-3 DNA plus various amounts of PSGL-1 DNA (1 to 100 ng) or PSGL-1 mutant DNA (1 to 500 ng). Virions were collected at 48 hours post-cotransfection and used to infect Rev-A3R5-GFP, using an equal p24 level of virus input. Virus infectivity was quantified with flow cytometry of HIV^+^ cells (GFP^+^) at 48 hours post infection. The 50% inhibitory dosage (IC_50_) was determined and plotted in (**C**). IC50 was plotted. The average IC_50_ values from multiple independent replicates (n = 4 to 10) were listed in (**D**). For comparison, vectors expressing APOBEC3G or SERINC5 were used and similarly quantified for effects on HIV-1 infectivity.

**Fig. 4.**
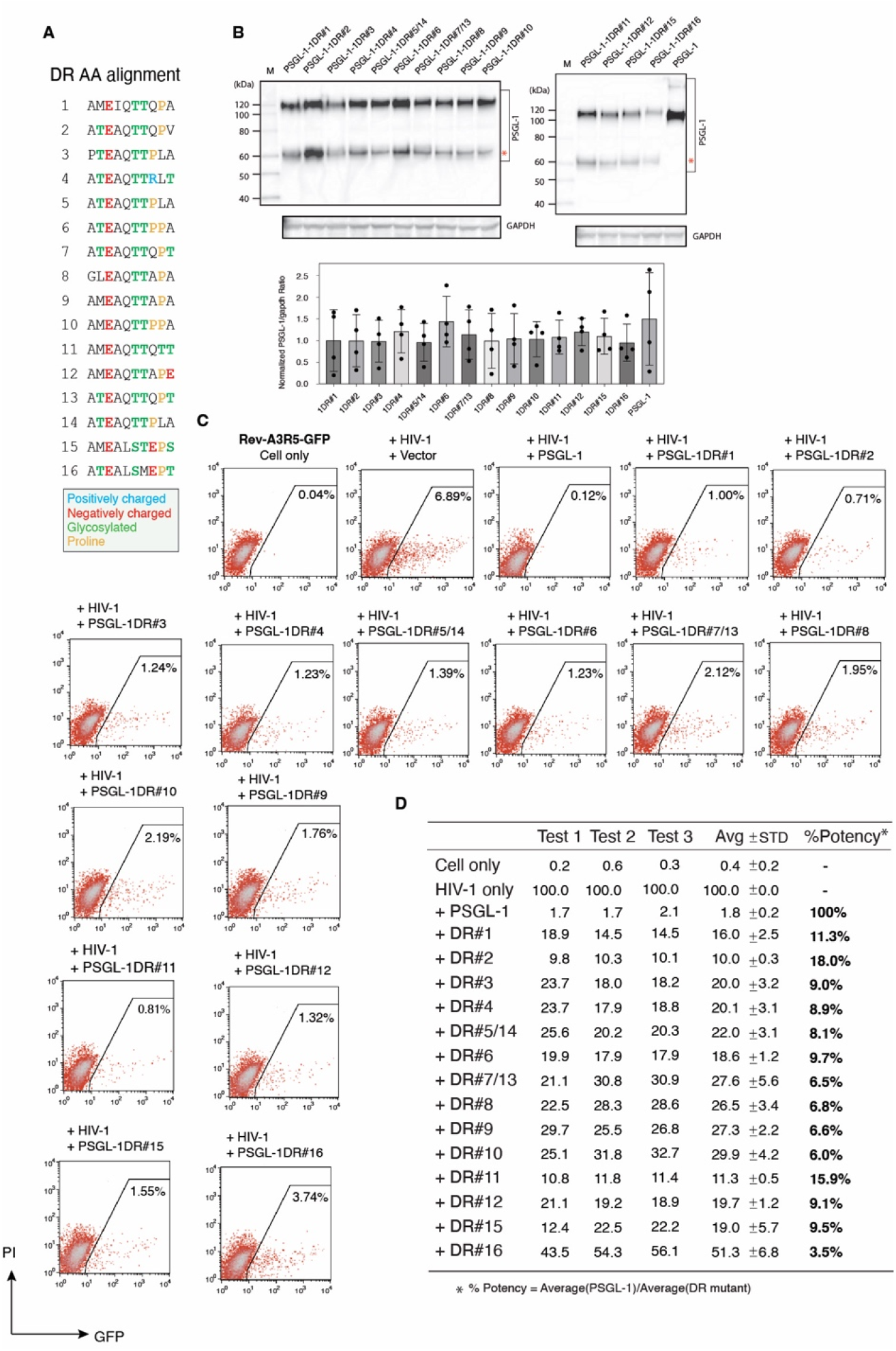
Quantification of the basal antiviral activities of individual single DRs. **(A)** Amino acid sequence alignment of the sixteen single DRs of PSGL-1. Among them, DR#5/14 and DR#7/13 have identical sequences. Charged and predicted glycosylated amino acids are labeled. **(B)** Western blot detection of PSGL-1 DR mutant proteins. HEK293T cells were transfected with 500 ng of each single DR mutants. Cell lysates were probed with an anti-PSGL-1 antibody or an anti-GAPDH antibody. The protein band intensity was quantified and the relative ratio of PSGL-1/GAPDH was calculated. The assays were performed in triplicate. Data are presented as mean ± SD of the three replicates. (**C**) Quantification of the relative antiviral activities of individual single DRs. HEK293T cells were co-transfected with 1 μg pNL4-3 DNA plus 200 ng PSGL-1 or each of the single-DR mutants DNA. An empty vector (+vector) was used to ensure the same amount of DNA was used in all cotransfections. Virions were collected at 48 hours post-transfection and used to infect Rev-A3R5-GFP indicator cells, using an equal level of p24 viral input. HIV-1 infectivity was quantified by flow cytometry. Results from three independent cotransfection and infection experiments were summarized in (**D**) and shown as the average plus standard derivation (Avg ± STD). “HIV-1+vector” was used as 100% infection. To calculate relative antiviral potency, “PSGL-1” was assigned as 100%, and the relative potency was calculated as “Average (PSGL-1)/Average (DR mutant %).

We also compared the anti-HIV activity of PSGL-1 with two other known HIV restriction factors, APOBEC3G (A3G) and SERINC5 (SER5) ^34-36^ (**Fig. 3D and Fig. S3**). Both proteins were expressed from the same CMV promotor as PSGL-1. Viral particles were similarly assembled in the presence of pcDNA3.1-A3G or pCMV6-SERINC5 (1 ng to 500 ng) and identically quantified in Rev-A3R5-GFP for IC_50_. A3G is potent against Vif-negative virus but less effective against the wild-type HIV-1 in the presence of Vif ^34^.

Consistently, in our assay, A3G was found to have a high IC_50_ of 355.2 ± 65.6 ng (n = 4) against HIV-1(NL4-3). In contrast, SERINC5 was found to have a low IC_50_ of 4.2 ± 1.7 ng (n = 4), a potency similar to PSGL-1 in restricting HIV-1. SERINC5 was also found to inhibit HIV virion release at high dosages (25 - 500 ng) (**Fig. S3**), and to inactivate HIV-1 infectivity at lower dosages (1 - 50 ng), which is different from PSGL-1. PSGL-1 does not block viral release even at a high dosage of 500 ng, and acts mainly through inactivation of virion infectivity ^17^.

### Quantification of the basal antiviral activities of individual single DRs

PSGL-1 DR contains 10 amino acids with the consensus sequence of “-A-T/M-E-A-Q-T-T-X-P/L-A/T-” which includes numerous O-glycosylated threonines (30%) and prolines (10%) (**Fig. 4A**). Given the variations between DR#1 and DR#16 in their antiviral activities (**Fig. 3D**), we were interested in quantifying and comparing the antiviral activities of all single DRs. Individual DRs were cloned into the pcDNA3.1 expression vector, and protein expression was confirmed by western blot (**Fig. 4B**). HIV-1 particles were assembled in the presence of 200 ng of each individual single DR mutant, and then quantified for virion infectivity using an equal p24 level of viral input (**Fig. 4C**). The experiments were independently repeated 3 times, and the relative percentages of GFP^+^ cells were converted into the relative antiviral potency (“PSGL-1” = 100%) (**Fig. 4D**). On average, the antiviral activity of individual single DR mutant varied from 3.5% (DR#16) to 18% (DR#2) of PSGL-1. These results suggest that a single DR in PSGL-1 can maintain basic antiviral activity. Among all the DRs, DR#16 exhibits the lowest antiviral activity (3.5%); however, its amino acid sequence shows greater divergence from the typical DR consensus sequence. The estimated cumulative potency of all 16 DRs can add to the full capacity of PSGL-1 (calculated at 143.5% of PSGL-1) (**Fig. 4D**).

### PSGL-1’s DR domain is required for blocking virion attachment to target cells

Mechanistically, the antiviral activity of PSGL-1 is mainly achieved through virion incorporation that sterically hinders particle attachment to target cells ^17^. We investigated whether DR is required for PSGL-1-mediated steric hindrance. Thus, we performed virion incorporation and attachment assays using particles produced in the presence of PSGL-1 or PSGL-1ΔDR. As shown in **Fig. 5A**, deleting all DRs (PSGL-1ΔDR) did not significantly affect virion incorporation of PSGL-1. Similarly, deleting PSGL-1 to a single DR (DR#1 and DR#16) also did not significantly affect their virion incorporation. However, when examined for the ability to block virion attachment to target cells, PSGL-1ΔDR lost the ability to block virion attachment (**Fig. 5B**). In addition, PSGL-1-DR#1 and PSGL-1-DR#16 also partially lost this ability. These results demonstrate that the DR domain is required for the steric hindrance activity of PSGL-1.

**Fig. 5.**
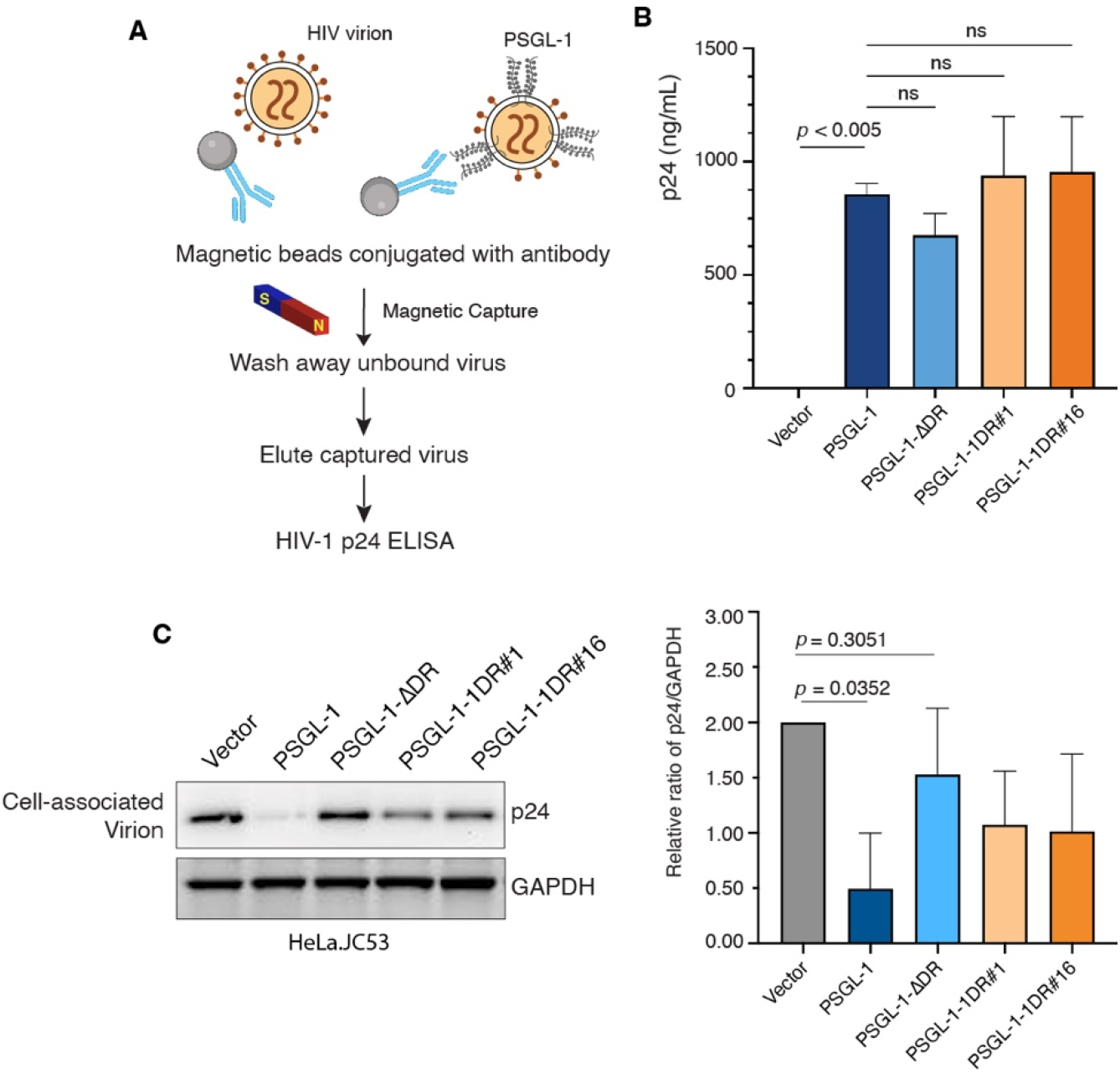
PSGL-1’s DR domain is required for blocking virion attachment to target cells. **(A)** Schematic of the immune-magnetic capture assay used to detect virion incorporation of PSGL-1 DR mutants. (**B**) Quantification of virion incorporation of PSGL-1 DR mutants. Shown are the ELISA p24 levels after capture. Results are from three independent replicates and presented as mean ± SD. *p*-values were based on three independent assays (ns, not significant, *p* > 0.05). **(C)** Quantification of virion attachment to target cells. Viral particles were produced by co-transfection of HEK293T cells with 1 μg pNL4-3 and 500 ng of PSGL-1, PSGL-1-ΔDR, PSGL-1 1DR#1, PSGL-1 1DR#16, or an empty vector. Viral particles were normalized by p24 levels and quantified for attachment to HeLa-JC.53 target cells via Western blot analysis of cell-bound p24. The protein band intensity was quantified and the relative ratios of p24/GAPDH were calculated. Vector p24/GAPDH ratios were assigned as “2”. The assays were performed in triplicate. Data are presented as mean ± SD of the three replicates.

### PSGL-1’s DR domain possesses antiviral activity independent of other PSGL-1 domains

Given the critical role of DR in PSGL-1’s antiviral activity, we investigated whether the DR domain alone possesses antiviral activity which can be independently transferred to other proteins. For this purpose, we inserted the whole DR domain (16 DRs) or a DR trimer (DR#14 to 16) into the extracellular domain of a T cell surface protein, CD2 (**Fig. 6A**). CD2 has two Ig-like domains in its extracellular domain, and the molecule has been shown to carry no anti-HIV activity ^37^. We constructed the hybrid vectors pCD2-16DRs and pCD2-Tri-DRs, in which the mucin-like DR domain was inserted into one of the Ig-like domains of CD2. The vectors were co-transfected with HIV-1 DNA into HEK293T cells to assemble virion particles. For comparison, the parental vector, pCMV6-XL5-CD2, was also similarly co-transfected. To confirm that CD2 is incorporated into virions, we performed magnetic bead-based antibody pull-down assay (**Fig. 6B**), and confirmed virion incorporation of CD2 (**Fig. 6C**). We subsequently quantified virion infectivity on Rev-A3R5-GFP indicator cells, using an equal p24 level of virus input. As shown in **Fig. 6D**, both CD2-Tri-DRs and CD2-16DRs had significantly decreased virion infectivity, and the 16 DRs had a stronger inhibition of HIV-1 infectivity than the Tri-DRs. These results demonstrate that the DR domain alone possesses partial anti-HIV activity which can be transferred to other proteins.

**Fig. 6.**
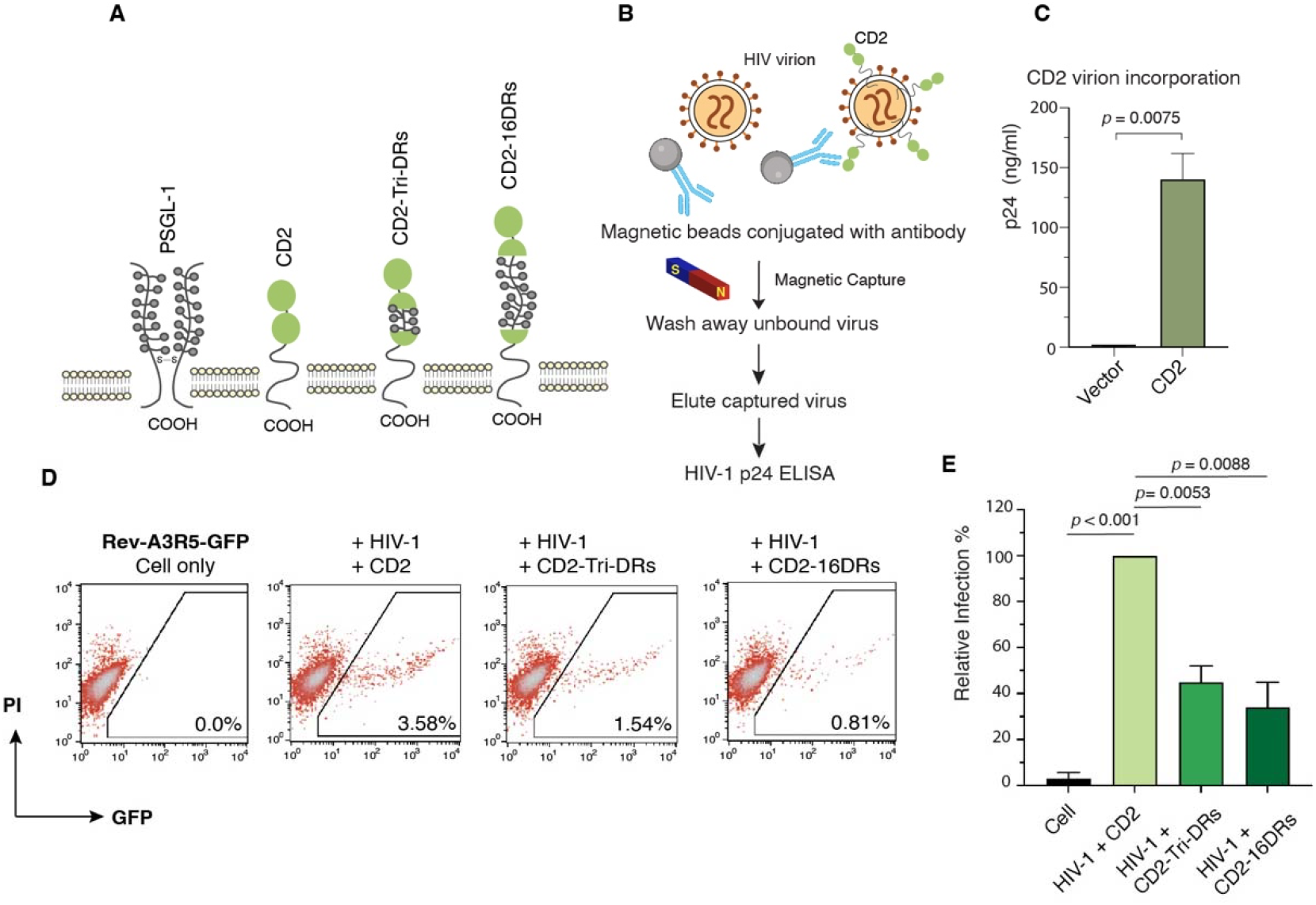
Chimeric CD2 with DR incorporation confers anti-HIV activity upon CD2. **(A)** Schematic of PSGL-1, CD2, CD2-TriDRs, and CD2-16DRs. (**B**) Schematic of the immune-magnetic capture assay used to detect CD2 on HIV-1 particles. (**C**) CD2 virion incorporation was confirmed in three independent experiments. (**D** and **E**) Inhibition of HIV infectivity by CD2-DR hybrid molecules. Virion particles were assembled in HEK293T cells by co-transfected with 1 μg pNL4-3 plus 500 ng the vector DNA (pCMV6-XL5-CD2, pCMV-CD2-TriDR, or pCMV-CD2-16DR). Virions were harvested at 48 hours post-transfection. Viral infectivity was quantified by infecting Rev-A3R5-GFP indicator cells with normalized p24 input. Relative infectivity assays were performed in triplicate. Data are presented as mean ± SD of the three replicates (**E**).

### Molecular modeling of DR’s antiviral activity

Given the variability (3.5% to 18%) in the anti-HIV potency of individual single DRs (**Fig. 4**), we were interested in knowing the molecular basis that determines DR’s antiviral potency. We performed molecular modeling using REST (Replica Exchange with Solute Tempering) simulations and evaluated the conformational ensembles of 14 single DR domains listed in **Fig. 7A**. To investigate potential structural determinants of DR potency, we computed the structural quantities as described in **Methods**. Among them were the distributions of DR backbone dihedral angles, *φ* and *ψ*, and the radial distribution functions *g*(*r*), measuring the number density of peptide atoms as a function of distance *r* to the peptide center of mass. Using *g*(*r*), we computed the locations of number density maxima *r*_*max*_ for each DR. **Fig. 7B** presents the average values of the structural quantities for each DR. The relationships between the two structural quantities and the antiviral potencies listed in **Fig. 7B** are probed in **Fig. 7C and 7D**. It is observed that the average dihedral angle ⟨*φ*⟩ and the location of peptide density peak *r*_*max*_ are likely to correlate with the potency. At the same time, no clear correlation is observed between ⟨*ψ*⟩ and antiviral potency. Indeed, the correlation coefficient *ρ* computed between the potency and ⟨*φ*⟩ is -0.59, but vanishes to 0.09 for ⟨*ψ*⟩. The correlation between *r*_*max*_ and potency is even greater with ρ being -0.64. It is important to mention that we have analyzed many other structural quantities pertaining to DRs, but none of them has ρ exceeding 0.5 in magnitude.

**Fig. 7.**
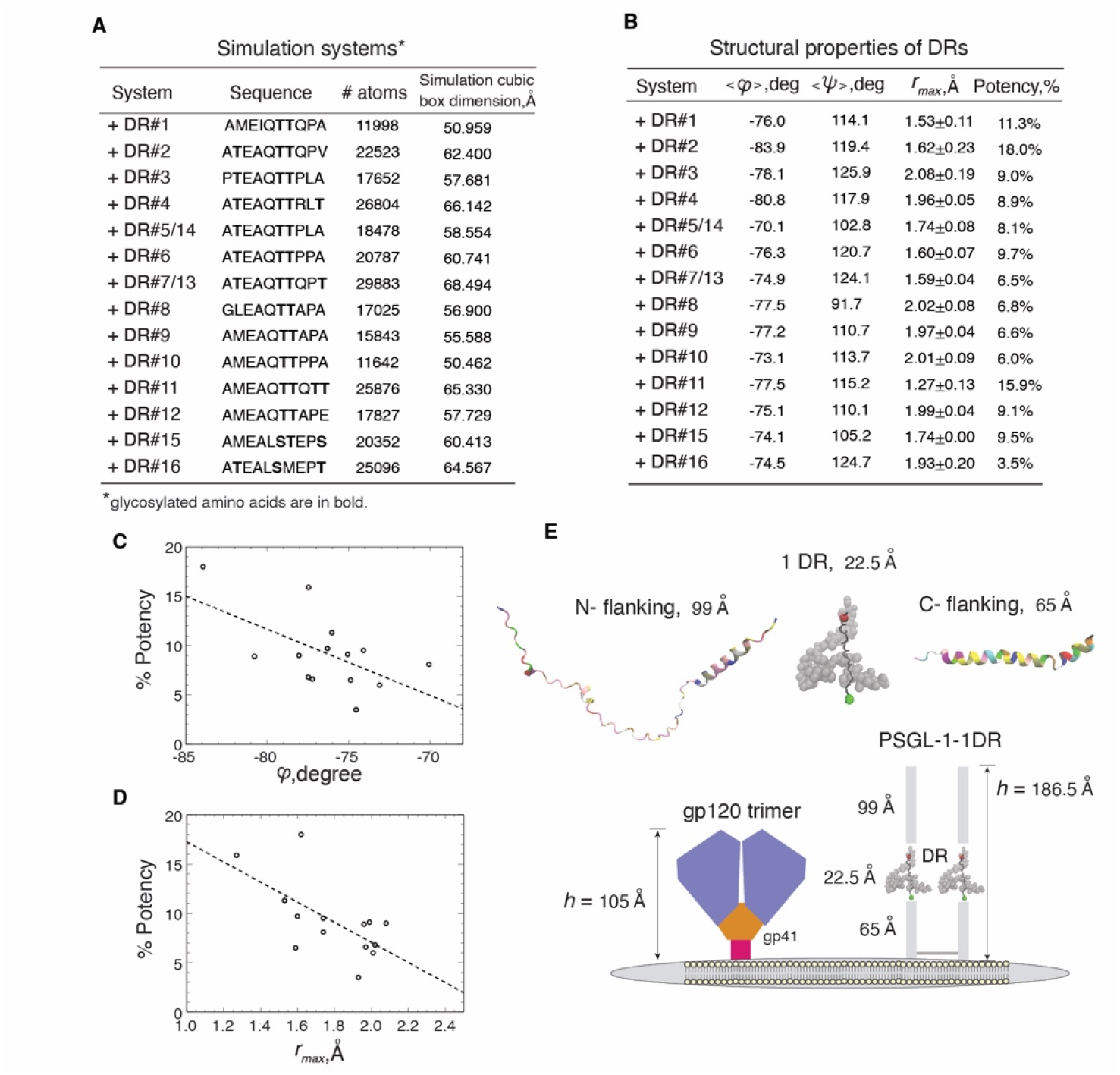
Molecular modeling of DR’s anti-HIV activity. **(A)** List of simulation systems. Shown are DR amino acid sequences, the number of atoms, and the cubic simulation of box dimension. (**B**) Summary of structural properties of DR. Shown are the average dihedral angle, ⟨*φ*⟩ and ⟨*ψ*⟩, as well as the location of peptide density peak r_max_ (± standard error). (**C** and **D**) Scatter plots illustrate the relationship between the potency against HIV infection and the average DR dihedral angle ⟨*φ*⟩ (**C**), or the location of the maximum peptide number density r_max_ (**D**). Dashed lines represent linear regression fits. (**E)** Hypothetical model of a single DR PSGL-1’s height. On the virion particle, PSGL-1-1DR mutant can pose as a fully extended “rod-like” structure. The height *h* of the single DR is measured by their end-to-end distance at 22.5 Å (the glycan side chains are shown in grey). The lengths of the N- and C-terminal flanking regions were modeled using AlphaFold with an end-to-end distance of 99 Å and 65 Å, respectively. The structural information of Env is based on published information.

What is a possible structural interpretation of the observed correlations? **Fig. 7B** shows that the average backbone dihedral angles ⟨*φ*⟩ and ⟨*ψ*⟩ occupy the upper left quadrant of Ramachandran plot, suggesting a prevalence of strand-like conformations. Indeed, clustering of DR peptide conformations shows that many DRs feature a single cluster with extended backbone (see **Fig. S5**). Furthermore, the average distance between the first and last C*φ* atoms *r*_*1N*_ in DRs ranges from 20 to 25 Å with the mean value of 22.5 Å. Since the average distance between adjacent C*φ* atoms in β-structure is 3.5 Å, *r*_*1N*_ of a DR adopting this conformation would be about 31.5 Å. Then, on average, DR backbones are extended from 63 to 79% of the maximum β-strand extension with the average extension across all DRs being 71%. It is noteworthy that this extension occurs without support of β-sheet structure but is driven by DR sequence properties and heavy glycosylation of DR sequences. In support of this conclusion, we found that compared to glycosylated DRs, the average *r*_*1N*_ for glycan-free DRs is reduced by about 10% to 20.8 Å. Then, the negative correlation between ⟨*φ*⟩ and potency implies that further extension of DR backbone, which is associated with a decrease in ⟨*φ*⟩ and better “compensation” of *ψ* angles, promotes DR potency. We also observed a strong correlation between the location of peptide density peak *r*_*max*_ and potency. **Fig. 7B** and **Fig. S4** reveal that between different DRs, *r*_*max*_ varies within about 2Å. Since the molecular weight of DR amino acids (excluding glycans) is fairly constant and the DRs predominantly sample extended conformations, the variations in *r*_*max*_ can be primarily attributed to the offsets of peptide center of mass from DR or to its bending. The latter is illustrated in **Fig. S5**. To test this interpretation of *r*_*max*_, we directly computed the distance between the peptide center of mass and the nearest peptide atom. This distance correlates with potency with ρ = -0.58 that is close to the correlation seen for *r*_*max*_. Thus, the negative correlation between *r*_*max*_ and potency indicates that the smaller the offset is and, consequently, the peptide bending, the higher potency becomes. Both observed correlations, involving ⟨*φ*⟩ and *r*_*max*_, point to the importance of extended DR backbone structure for antiviral potency.

## DISCUSSIONS

In this study, we investigated and compared the relative antiviral activities of DR mutants and discovered several key findings: 1. Deleting all DRs significantly diminished PSGL-1’s anti-HIV activity (IC_50_>400 ng), highlighting the essential role of DR in PSGL-1’s antiviral action. 2. DR mutants with individual single DRs exhibited basal anti-HIV activity, ranging from 3.5% to 18% of the full-length PSGL-1. 3. Individual DRs have slightly varied sequences (with the exception of DR#5/14 and DR#7/13) and differing anti-HIV potency. 4. The anti-HIV activity of DR appears to be cumulative; an increase in the number of DRs correlates with enhanced antiviral potency. 5. Molecular modeling revealed a structure-function relationship between the anti-HIV potency of DRs and the bending of the DR backbone. These findings suggest that DR acts as a crucial antiviral unit in PSGL-1’s restriction mechanism. Nevertheless, we note that the number of natural DR sequences analyzed in this study is limited. Hence, the proposed structural-function relationship should be regarded as constrained, and future studies utilizing machine learning on larger datasets of artificially designed DR variants are necessary for more precise molecular predictions.

Mechanistically, PSGL-1 inhibits HIV infectivity through two distinct mechanisms: 1. Steric hindrance, a mechanism by which PSGL-1 is incorporated into virion, leading to steric hinderance of virion attachment to target cells ^17,18^; 2. Spatial hindrance, a mechanism by which PSGL-1 excludes HIV Env incorporation into virion particles ^17,38^. In both mechanisms, DR is vital ^38^ (**Fig. 5**). Notably, excluding Env requires a minimum of nine DRs ^38^, while, in this study, a single DR was found to suffice to maintain basal anti-HIV activity (**Fig. 2** and **3**). Molecular modeling indicates that PSGL-1 mutants with seven or fewer DRs assume as extended “rod-like” structure, while those with nine or more DRs adopt a “coil-like” structure, likely enhancing their ability to spatially exclude Env ^38^. For PSGL-1, preventing virion attachment through steric hindrance appears to be the dominant phenotype compared to its capacity for Env exclusion ^17^, with different minimal DR requirements for each mechanism.

Our molecular simulations imply that the extension of DR is critical for antiviral activity (**Fig. 7**). It is reasonable to hypothesize that PSGL-1 mutants with a single DR would be elongated enough to interfere with Env binding to target cells. Based on prior studies, the Env trimer’s height (*h)* is around 105 Å on virion membrane ^39-41^, while the estimated height of fully-extended extracellular domain of a single DR mutant is at approximately 187 Å (**Fig. 7E**), likely sufficient for basal steric hindrance.

We also observed that the antiviral potency of individual DRs within full-length PSGL-1 is not randomly distributed along the PSGL-1 backbone (**Fig. 7B** and **Fig. S6**). In the DR domain (excluding DR#16), DRs with stronger antiviral activities are located at both the top (DR#1 to #6, average potency 10.8%) and the bottom (DR#11 to #15, average potency 9.8%), while those with weaker activities are found in the middle (DR#7 to #10, average potency 6.4%). The reason for this non-random distribution remains unclear.

Given that a DR’s antiviral potency is associated with its structural extension, it is reasonable to speculate that the observed differences in antiviral efficacy may reflect DR’s molecular rigidity; the middle section (DR#7 to #10) might be more bended than the top or bottom ones (**Fig. S6**). Indeed, the average *r*_*max*_ values measuring DR bending for the top and bottom sections are 1.76±0.06 and 1.71±0.02 Å, whereas the middle section features *r*_*max*_ at 1.90±0.01 Å. It is also conspicuous that the middle section of DRs (#7 to #10) corresponds to the onset of extension-collapse transition in DR tandems that lends further support for enhanced bending propensity harbored in this PSGL-1 section ^38^.

We demonstrated that the intrinsic anti-HIV activity of DR is transferrable independent of the other PSGL-1 domains; inserting the DR domain into the extracellular Ig-like domain of CD2 permitted the hybrid CD2-DR molecules to partially acquire the anti-HIV properties of PSGL-1 (**Fig. 6**). However, the hybrid CD2-DR molecules are not as potent as PSGL-1 for inhibiting HIV-1. A possible reason could be that the lack of PSGL-1’s intracellular domain in the hybrid CD2-DR molecules may decrease the efficiency of virion incorporation. The polybasic sequence in PSGL-1’s cytoplasmic domain has been shown to selectively direct PSGL-1 to the site of HIV virion assembly ^42,43^. Although CD2 was found to be incorporated into virion (**Fig. 6B**), there is no evidence that CD2 co-clusters with HIV Gag at the virion assembly site for enriched selective incorporation ^44,45^.

## Acknowledgments

The authors wish to thank the NIH AIDS Reagent Program for reagents. This work was supported by Public Health Service grant R01AI148012 (to Y.W.) and R56AI183995 (to Y.W., S.J., and D.K.K).

## Author contributions

Y.W. conceived and supervised the study. Y.W. performed experimental data analysis.

D.K. and M.S.J. supervised molecular modeling. B.M.D., C.L., D.K., and M.S.J. performed molecular modeling. Y.H., L.S., Y.F., L.C., and S.T. performed experiments.

Y.W. wrote the original manuscript draft and D.K. wrote the molecular modeling section. All authors reviewed and edited the manuscript.

## Competing interests

Patents related to PSGL-1 have been filed by George Mason University.

**Correspondence and requests for materials should be addressed to Y.W**

## References

1 Sako, D. et al. Expression cloning of a functional glycoprotein ligand for P-selectin. Cell 75, 1179–1186 (1993).

2 Laszik, Z. et al. P-selectin glycoprotein ligand-1 is broadly expressed in cells of myeloid, lymphoid, and dendritic lineage and in some nonhematopoietic cells. Blood 88, 3010–3021 (1996).

3 Fujimoto, T. T. et al. Expression and functional characterization of the P-selectin glycoprotein ligand-1 in various cells. Int J Hematol 64, 231–239 (1996).

4 Almulki, L. et al. Surprising up-regulation of P-selectin glycoprotein ligand-1 (PSGL-1) in endotoxin-induced uveitis. FASEB J 23, 929–939 (2009). 10.1096/fj.08-118760

5 Xu, H. et al. Recruitment of IFN-gamma-producing (Th1-like) cells into the inflamed retina in vivo is preferentially regulated by P-selectin glycoprotein ligand 1:P/E-selectin interactions. J Immunol 172, 3215–3224 (2004).

6 Schumacher, A. et al. P-selectin glycoprotein ligand-1 (PSGL-1) is up-regulated on leucocytes from patients with chronic obstructive pulmonary disease. Clin Exp Immunol 142, 370–376 (2005). 10.1111/j.1365-2249.2005.02920.x

7 Somers, W. S., Tang, J., Shaw, G. D. & Camphausen, R. T. Insights into the molecular basis of leukocyte tethering and rolling revealed by structures of P- and E-selectin bound to SLe(X) and PSGL-1. Cell 103, 467–479 (2000).

8 Li, F. et al. Visualization of P-selectin glycoprotein ligand-1 as a highly extended molecule and mapping of protein epitopes for monoclonal antibodies. J Biol Chem 271, 6342–6348 (1996).

9 Patel, K. D., Nollert, M. U. & McEver, R. P. P-selectin must extend a sufficient length from the plasma membrane to mediate rolling of neutrophils. J Cell Biol 131, 1893–1902 (1995).

10 McEver, R. P., Moore, K. L. & Cummings, R. D. Leukocyte trafficking mediated by selectin-carbohydrate interactions. J Biol Chem 270, 11025–11028 (1995).

11 Li, F. et al. Post-translational modifications of recombinant P-selectin glycoprotein ligand-1 required for binding to P- and E-selectin. J Biol Chem 271, 3255–3264 (1996).

12 Fang, Y., Wu, J., McEver, R. P. & Zhu, C. Bending rigidities of cell surface molecules P-selectin and PSGL-1. Journal of Biomechanics 42, 303–307 (2009). 10.1016/j.jbiomech.2008.11.020

13 Umeki, S. et al. Anti-adhesive property of P-selectin glycoprotein ligand-1 (PSGL-1) due to steric hindrance effect. J Cell Biochem 114, 1271–1285 (2013). 10.1002/jcb.24468

14 Afshar-Kharghan, V., Diz-Kucukkaya, R., Ludwig, E. H., Marian, A. J. & Lopez, J. A. Human polymorphism of P-selectin glycoprotein ligand 1 attributable to variable numbers of tandem decameric repeats in the mucinlike region. Blood 97, 3306–3307 (2001).

15 Tauxe, C. et al. P-selectin glycoprotein ligand-1 decameric repeats regulate selectin-dependent rolling under flow conditions. J Biol Chem 283, 28536–28545 (2008). 10.1074/jbc.M802865200

16 Liu, Y. et al. Proteomic profiling of HIV-1 infection of human CD4(+) T cells identifies PSGL-1 as an HIV restriction factor. Nat Microbiol 4, 813–825 (2019). 10.1038/s41564-019-0372-2

17 Fu, Y. et al. PSGL-1 restricts HIV-1 infectivity by blocking virus particle attachment to target cells. Proc Natl Acad Sci U S A 117, 9537–9545 (2020). 10.1073/pnas.1916054117

18 Murakami, T., Carmona, N. & Ono, A. Virion-incorporated PSGL-1 and CD43 inhibit both cell-free infection and transinfection of HIV-1 by preventing virus-cell binding. Proc Natl Acad Sci U S A 117, 8055–8063 (2020). 10.1073/pnas.1916055117

19 Huang, J. et al. CHARMM36m: an improved force field for folded and intrinsically disordered proteins. Nature Methods 14, 71–73 (2017). 10.1038/nmeth.4067

20 Guvench, O. et al. CHARMM Additive All-Atom Force Field for Carbohydrate Derivatives and Its Utility in Polysaccharide and Carbohydrate–Protein Modeling. Journal of Chemical Theory and Computation 7, 3162–3180 (2011). 10.1021/ct200328p

21 Cummings, R. D. Structure and function of the selectin ligand PSGL-1. Braz J Med Biol Res 32, 519–528 (1999). 10.1590/s0100-879×1999000500004

22 Jorgensen, W. L., Chandrasekhar, J., Madura, J. D., Impey, R. W. & Klein, M. L. Comparison of simple potential functions for simulating liquid water. The Journal of Chemical Physics 79, 926–935 (1983). 10.1063/1.445869

23 MacKerell, A. D., Jr. et al. All-Atom Empirical Potential for Molecular Modeling and Dynamics Studies of Proteins. The Journal of Physical Chemistry B 102, 3586–3616 (1998). 10.1021/jp973084f

24 Wang, L., Friesner, R. A. & Berne, B. J. Replica exchange with solute scaling: a more efficient version of replica exchange with solute tempering (REST2). J Phys Chem B 115, 9431–9438 (2011). 10.1021/jp204407d

25 Sugita, Y. & Okamoto, Y. Replica-exchange molecular dynamics method for protein folding. Chemical Physics Letters 314, 141–151 (1999). 10.1016/S0009-2614(99)01123-9

26 Smith, A. K., Lockhart, C. & Klimov, D. K. Does Replica Exchange with Solute Tempering Efficiently Sample Aβ Peptide Conformational Ensembles? Journal of Chemical Theory and Computation 12, 5201–5214 (2016). 10.1021/acs.jctc.6b00660

27 Liu, P., Kim, B., Friesner, R. A. & Berne, B. J. Replica exchange with solute tempering: A method for sampling biological systems in explicit water. Proceedings of the National Academy of Sciences 102, 13749–13754 (2005). doi:10.1073/pnas.0506346102

28 Best, R. B. et al. Optimization of the Additive CHARMM All-Atom Protein Force Field Targeting Improved Sampling of the Backbone □, ψ and Side-Chain χ1 and χ2 Dihedral Angles. Journal of Chemical Theory and Computation 8, 3257–3273 (2012). 10.1021/ct300400x

29 Zhang, Y., Liu, X. & Chen, J. Re-Balancing Replica Exchange with Solute Tempering for Sampling Dynamic Protein Conformations. Journal of Chemical Theory and Computation 19, 1602–1614 (2023). 10.1021/acs.jctc.2c01139

30 Humphrey, W., Dalke, A. & Schulten, K. VMD: Visual molecular dynamics. Journal of Molecular Graphics 14, 33–38 (1996). 10.1016/0263-7855(96)00018-5

31 Daura, X. et al. Peptide Folding: When Simulation Meets Experiment. Angewandte Chemie International Edition 38, 236–240 (1999). 10.1002/(SICI)1521-3773(19990115)38:1/2<236::AID-ANIE236>3.0.CO;2-M

32 Wu, Y., Beddall, M. H. & Marsh, J. W. Rev-dependent indicator T cell line. Current HIV Research 5, 395–403 (2007).

33 Wu, Y., Beddall, M. H. & Marsh, J. W. Rev-dependent lentiviral expression vector. Retrovirology 4, 12 (2007).

34 Sheehy, A. M., Gaddis, N. C., Choi, J. D. & Malim, M. H. Isolation of a human gene that inhibits HIV-1 infection and is suppressed by the viral Vif protein. Nature 418, 646–650 (2002).

35 Usami, Y., Wu, Y. & Gottlinger, H. G. SERINC3 and SERINC5 restrict HIV-1 infectivity and are counteracted by Nef. Nature 526, 218–223 (2015). 10.1038/nature15400

36 Rosa, A. et al. HIV-1 Nef promotes infection by excluding SERINC5 from virion incorporation. Nature 526, 212–217 (2015). 10.1038/nature15399

37 Dabbagh, D. et al. Identification of the SHREK Family of Proteins as Broad-Spectrum Host Antiviral Factors. Viruses 13, 832 (2021). 10.3390/v13050832

38 Tiwari, S. et al. PSGL-1 excludes HIV Env from virion surface through spatial hindrance involving structural folding of the decameric repeats (DR). bioRxiv, 2024.2012.2028.630612 (2024). 10.1101/2024.12.28.630612

39 Li, Z. et al. Subnanometer structures of HIV-1 envelope trimers on aldrithiol-2-inactivated virus particles. Nature Structural & Molecular Biology 27, 726–734 (2020). 10.1038/s41594-020-0452-2

40 Liu, J., Bartesaghi, A., Borgnia, M. J., Sapiro, G. & Subramaniam, S. Molecular architecture of native HIV-1 gp120 trimers. Nature 455, 109–113 (2008). 10.1038/nature07159

41 Zhu, P. et al. Distribution and three-dimensional structure of AIDS virus envelope spikes. Nature 441, 847–852 (2006).

42 Grover, J. R., Veatch, S. L. & Ono, A. Basic motifs target PSGL-1, CD43, and CD44 to plasma membrane sites where HIV-1 assembles. J Virol 89, 454–467 (2015). 10.1128/JVI.02178-14

43 Llewellyn, G. N., Grover, J. R., Olety, B. & Ono, A. HIV-1 Gag associates with specific uropod-directed microdomains in a manner dependent on its MA highly basic region. J Virol 87, 6441–6454 (2013). 10.1128/JVI.00040-13

44 Burnie, J. et al. Identification of CD38, CD97, and CD278 on the HIV surface using a novel flow virometry screening assay. Scientific Reports 13, 23025 (2023). 10.1038/s41598-023-50365-0

45 Burnie, J. & Guzzo, C. The Incorporation of Host Proteins into the External HIV-1 Envelope. Viruses 11 (2019). 10.3390/v11010085

