## Supplementa Figures 1-6 and Tables 1-2 for "The decameric repeat (DR) of PSGL-1 functions as a basic antiviral unit in restricting HIV-1 infectivity"

### Supplementary Information

#### Molecular Modeling

**Equilibration of DR conformations:** To evaluate DR equilibration in REST simulations, we partitioned simulation time in 30 ns batches. For each batch we computed the correlation coefficient $\rho$ between the potency and the location of peptide number density maximum $r_{max}$. The resulting data are shown in **Table S1**. We found that $\rho$ increases with time till 60 ns and then levels off for the last two batches. Based on this observation, we assumed that for all DRs the last 60 ns of sampling is equilibrated.

**Correlation cross-validation:** To ensure that our correlation analysis of DR properties is robust, we performed leave-one-out cross-validation by consecutively removing individual DRs from the dataset and recomputing Pearson's correlation coefficient $\rho$ with potency. When cross-validation was performed for $\langle\varphi\rangle$, the largest change in $\rho$ occurs when DR#2 is removed, resulting in $\rho$ = -0.27. For comparison, $\langle\psi\rangle$ consistently shows weak correlations in the cross-validation analysis. In contrast, for $r_{max}$ $\rho$ remains consistent, with DR#11 being the biggest outlier resulting in $\rho$ = -0.48. These findings are confirmed by visual inspection of respective scatter plots in **Fig. S4**. Because the correlation of $r_{max}$ with potency is cross-validated and it is structurally linked to the average dihedral angle $\langle\varphi\rangle$, we retained both structural measures in our analysis.

**Distributions of peptide number densities:** The radial distribution functions $g(r)$ measuring the number density of peptide atoms as a function of distance $r$ to the peptide center of mass were computed for all DRs. The examples of $g(r)$ illustrating their variability are shown in **Fig. S4.**

**DR clustering:** Following the procedure described in Methods, we performed clustering of DR conformations. The fractions of DR conformations collected in the largest cluster range from 0.19 to 0.63 implicating a generally rigid DR peptide structure. The centroids of the largest clusters for DR#3 and #11 are shown in **Fig. S5**. These DRs were selected because they exhibit the largest variation in the location of peptide number density $r_{max}$. Consistent with our structural interpretation of $r_{max}$, DR#11, which features the smallest $r_{max}$, adopts an extended conformation with minimal bending, whereas DR#3 featuring the largest $r_{max}$ exhibits a noticeable bend.

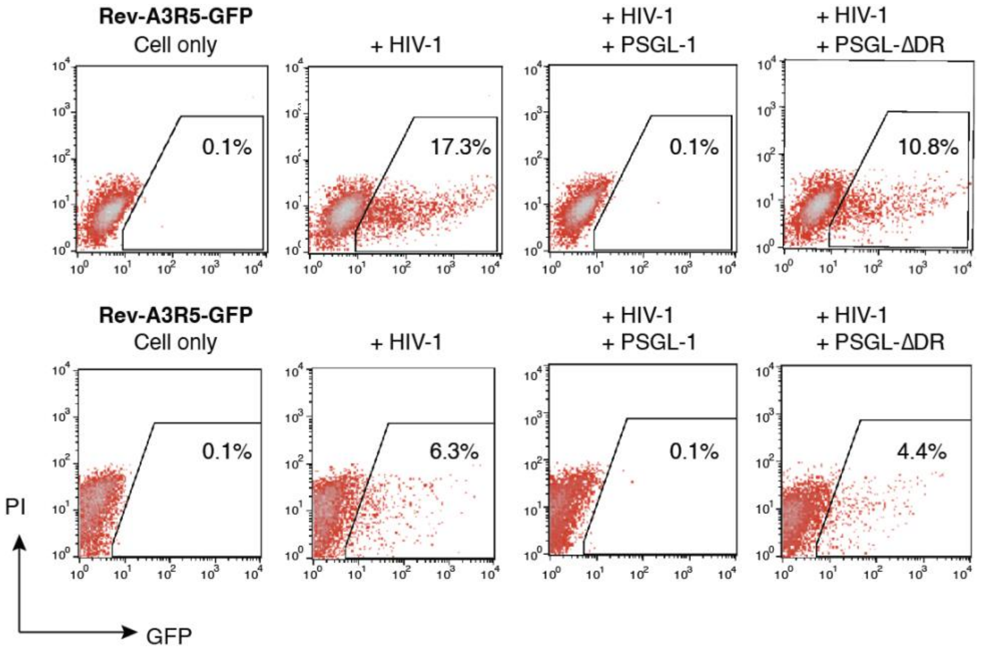


**Fig. S1. PSGL-1's DR domain is required for its antiviral activity.**
Quantification of the antiviral activities of the DR deletion mutant. HEK293T cells were co-transfected with 1 µg pNL4-3 DNA plus 500 ng PSGL-1 or PSGL-1ΔDR, or an empty vector. Virions were harvested at 48 hours post-transfection and used to infect Rev-A3R5-GFP indicator cells, using an equal p24 level of virus input. Virus infectivity was quantified with flow cytometry of $HIV^+$ cells ($GFP^+$). PI (propidium iodide) was used to examine GFP only in viable cells.

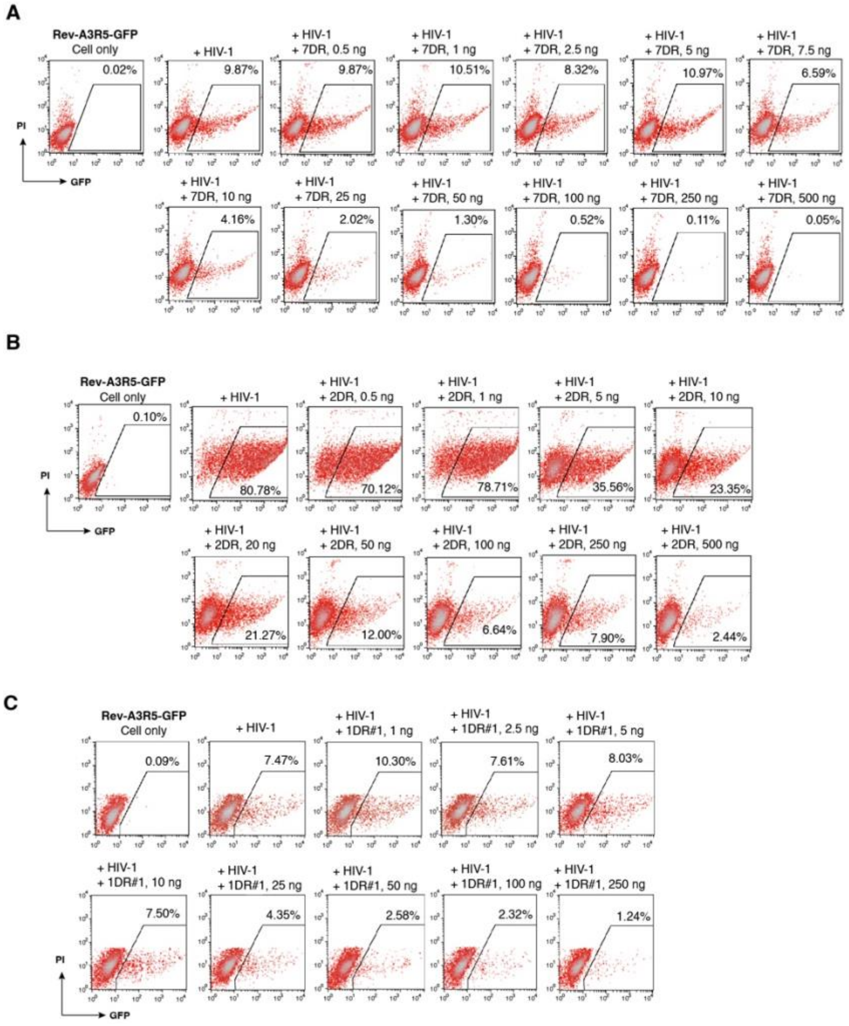


**Fig.S2. Quantification of the 50% inhibitory dosage ($IC_{50}$) of PSGL-1 and its DR deletion mutants using the HIV Rev-dependent Rev-A3R5-GFP cells**. HEK293T cells were cotransfected with 1 µg pNL4-3 DNA plus various amounts of PSGL-1 mutant DNA (1 to 250 or 500 ng). Virions were collected at 48 hours post-cotransfection and used to infect Rev-A3R5-GFP, using an equal p24 level of virus input. Virus infectivity was quantified with flow cytometry of $HIV^+$ cells ($GFP^+$) at 48 or 72 hours post infection. Shown are $IC_{50}$ quantifications for PSGL-1-7DR (**A**), PSGL-1-2DR (**B**), and PSGL-1DR#1 (**C**).

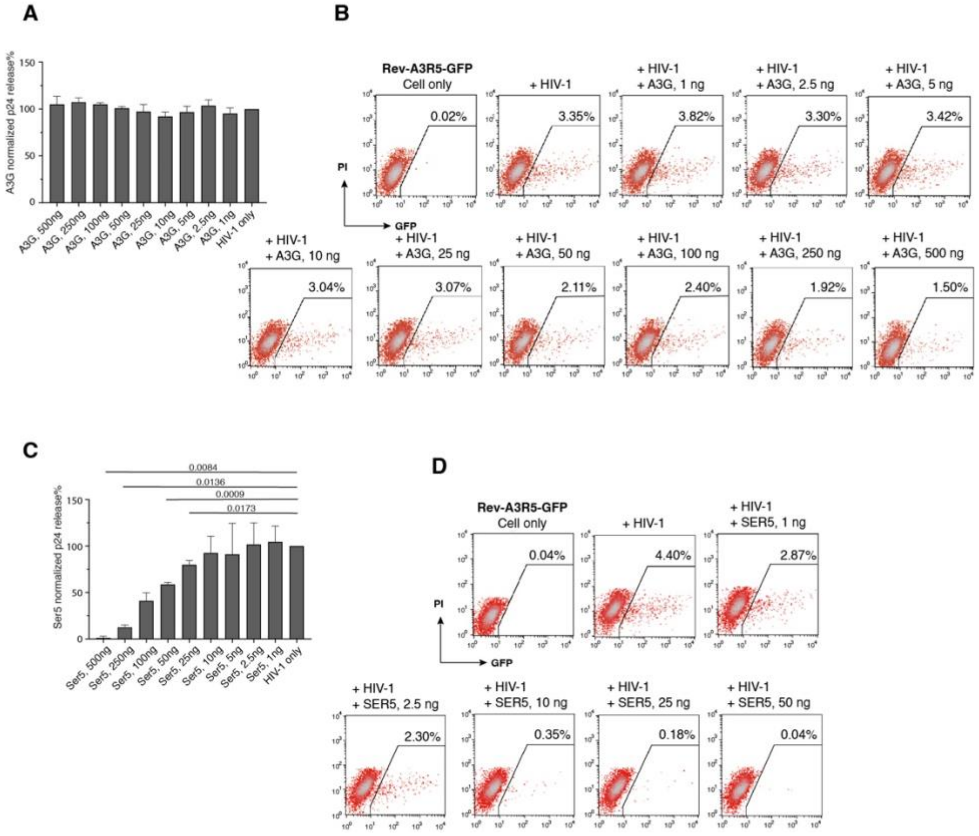


**Fig.S3. Quantification and comparison of HIV restriction factor APOBEC3G (A3G) or SERINC5**. **(A)** Effects of APOBEC3G on HIV-1 virion release. HEK293T cells were co-transfected with HIV-1 DNA (1 µg) plus an A3G-expression vector (1 to 500 ng of DNA). Virion release was quantified at 48 hours post co-transfection by HIV-1 p24 ELISA. Data are represented as mean ± SD from assay triplicate. (**B**) Quantification of the 50% inhibitory dosage ($IC_{50}$) of A3G using the HIV Rev-dependent Rev-A3R5-GFP cells. HEK293T cells were cotransfected with 1 µg pNL4-3 DNA plus various amounts of A3G DNA (1 to 500 ng). Virions were collected at 48 hours post-cotransfection and used to infect Rev-A3R5-GFP, using an equal p24 level of virus input. Virus infectivity was quantified with flow cytometry of $HIV^+$ cells ($GFP^+$) at 48 hours post infection. **(C)** Effects of SERINC5 on HIV-1 virion release. HEK293T cells were co-transfected with HIV-1 DNA (1 µg) plus a SERINC5-expression vector (1 to 500 ng of DNA). Virion release was quantified at 48 hours post co-transfection by HIV-1 p24 ELISA. Data are represented as mean ± SD from assay triplicate. (**D**) Quantification of the 50% inhibitory dosage ($IC_{50}$) of SERINC5 using the HIV Rev-dependent Rev-A3R5-GFP cells. HEK293T cells were cotransfected with 1 µg pNL4-3 DNA plus various amounts of SER5 DNA (1 to 50 ng). Virions were collected at 48 hours post-cotransfection and used to infect Rev-A3R5-GFP, using an equal p24 level of virus input. Virus infectivity was quantified with flow cytometry of $HIV^+$ cells ($GFP^+$) at 48 hours post infection.

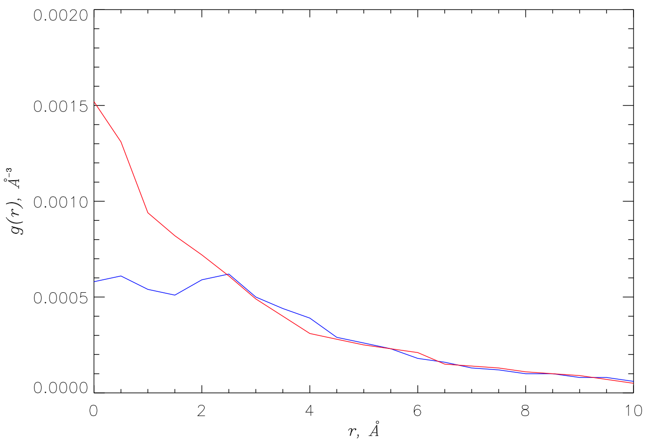


**Fig. S4** The radial distribution functions $g(r)$ measuring the number density of peptide atoms as a function of distance *r* to the peptide center of mass are plotted for DR#3 (blue) and #11 (red). These DRs feature the largest and the smallest $r_{max}$, the location of peptide density maximum. The figure reveals that the variations in $g(r)$ are confined to $r$ < 2.5 Å.

(a) (b)

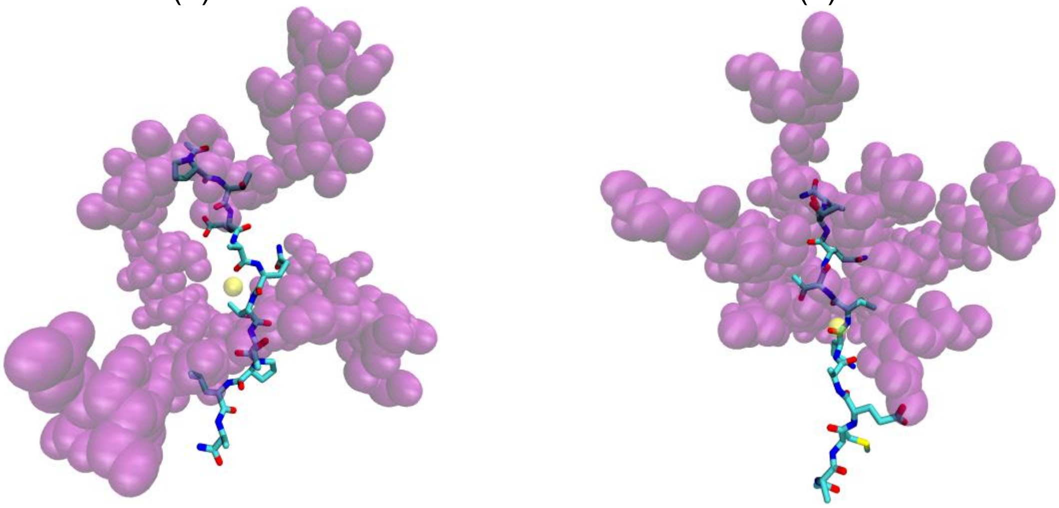

**Fig. S5** DR peptides are represented by their largest conformational clusters. Specifically, panels (a) and (b) present the centroids of the clusters with populations 63% and 31% for DR#3 and DR#11, respectively. The glycan side chains are shown in purple. The location of peptide center of mass is marked by a yellow sphere. DR#11 reveals minimal bending compared to DR#3.

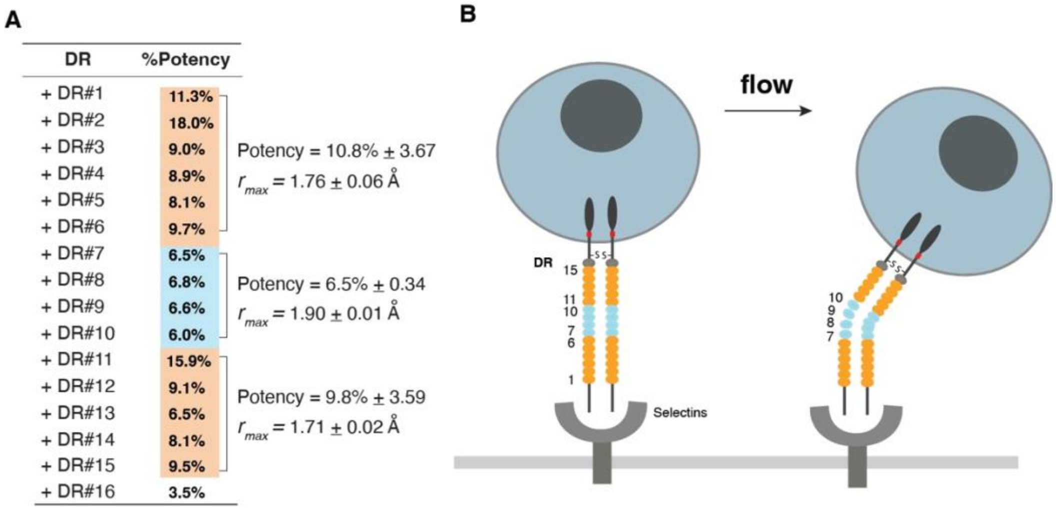

A

| DR | %Potency |
|---|---|
| + DR#1 | 11.3% |
| + DR#2 | 18.0% |
| + DR#3 | 9.0% |
| + DR#4 | 8.9% |
| + DR#5 | 8.1% |
| + DR#6 | 9.7% |
| + DR#7 | 6.5% |
| + DR#8 | 6.8% |
| + DR#9 | 6.6% |
| + DR#10 | 6.0% |
| + DR#11 | 15.9% |
| + DR#12 | 9.1% |
| + DR#13 | 6.5% |
| + DR#14 | 8.1% |
| + DR#15 | 9.5% |
| + DR#16 | 3.5% |

Potency = 10.8% ± 3.67, $r_{max}$ = 1.76 ± 0.06 Å (DR#1–#6)

Potency = 6.5% ± 0.34, $r_{max}$ = 1.90 ± 0.01 Å (DR#7–#10)

Potency = 9.8% ± 3.59, $r_{max}$ = 1.71 ± 0.02 Å (DR#11–#15)



**Fig.S6. Hypothetical model of a possible connection between DR's antiviral potency and its structural flexibility. (A)** List of the relative antiviral potency of individual single DRs. (**B**) Hypothetical model, in which the weaker antiviral potency of DR#7-#10, situated in the middle of the DR domain, may reflect their structural flexibility for bending, facilitating PSGL-1-mediated rolling of lymphocytes on endothelium. The average potency and the average $r_{max}$ of the three sections of DR are shown.

**Table S1** Correlations between potency and the location of peptide number density peak within simulation batches.

| **Batch range *t*, ns** | $\rho$ |
|---|---|
| $t < 30$ | -0.36 |
| $30 < t < 60$ | -0.40 |
| $60 < t < 90$ | -0.56 |
| $90 < t < 120$ | -0.54 |

**Table S2** Correlation coefficients $\rho$ for structural quantities and potency observed upon leave-one-out cross-validation.

| Excluded DR | $\langle\varphi\rangle$ | $\langle\psi\rangle$ | $r_{max}$ |
|---|---|---|---|
| DR#1 | -0.60 | 0.09 | -0.63 |
| DR#2 | -0.27 | -0.03 | -0.69 |
| DR#3 | -0.60 | 0.10 | -0.68 |
| DR#4 | -0.64 | 0.09 | -0.65 |
| DR#5 | -0.65 | 0.06 | -0.65 |
| DR#6 | -0.59 | 0.08 | -0.65 |
| DR#7 | -0.48 | 0.15 | -0.72 |
| DR#8 | -0.62 | -0.05 | -0.62 |
| DR#9 | -0.61 | 0.07 | -0.62 |
| DR#10 | -0.56 | 0.09 | -0.61 |
| DR#11 | -0.63 | 0.08 | -0.48 |
| DR#12 | -0.59 | 0.09 | -0.66 |
| DR#15 | -0.60 | 0.10 | -0.64 |
| DR#16 | -0.58 | 0.26 | -0.64 |
